# clonotypeR—high throughput analysis of T cell antigen receptor sequences

**DOI:** 10.1101/028696

**Authors:** Charles Plessy, Encarnita Mariotti-Ferrandiz, Ri-Ichiroh Manabe, Shohei Hori

## Abstract

**Motivation:** The T cell receptors are expressed as millions of different rearrangements. Amplified as a complex mixture of PCR products, they can be sequenced directly on *next-generation* instruments without the need for cloning. This method is increasingly used to characterize, quantify and study these highly diverse receptors.

**Results:** We present here clonotypeR, a software package to identify and analyze antigen receptors from high-throughput sequence libraries. ClonotypeR is designed to process, organize and analyze very large numbers of sequences, in the order of millions, typically produced by Roche 454 or Illumina instruments, and is made of two parts. The first contains shell scripts and reference segment sequences to produce a data file where each line represents a the detection of a clonotype in a sequence read. The second part is a R module available from Bioconductor, to load and filter the data, and prepare clonotype abundance tables ready for analysis with third-party tools for differential representation analysis, sample clustering, etc. To analyze clonotype data at the nucleotide level, we introduce unique clonotype identifiers based on those developed by Yassai et al. (2009), that we corrected to avoid identifier collisions.

**Availability:** http://clonotyper.branchable.com (CC0 license).

## Introduction

The adaptive immune system cells recognize their targets by producing a large number of variable receptors, where the sequence diversity, not encoded directly in the germ line, originates through *somatic recombination*. Most T lymphocytes express only one receptor variant, and the collection of all the different receptors in a cell population is called a *repertoire*. Once the somatic recombination is done, the receptor is transmitted to daughter cells across divisions. The collection of cells expressing the same receptor, and by extension the receptor itself, is called a *clonotype*.

The somatic recombination assembles a mature gene from arrays of *segments* on the genome (Tonegawa, 1983, Chien et al., 1984). The junction between the N-terminal *V* and the downstream *J* segment, with the possible inclusion of a *D* segment, is called the *complementarity determining region 3* (CDR3) and is one of the key contact points with the antigen. The somatic recombination is imprecise and therefore the sequence of the CDR3 has to be determined experimentally. Flanking the CDR3, there are two structural motifs that are essential for the receptor’s function and therefore strongly conserved: a cysteine at the end of the V segment and a FGxG motif in the J segment (Sant’Angelo et al., 1997). Characterizing the receptor clonotype diversity requires the identification of the V and J segments and sequencing the CDR3, as well as suitable and robust statistics (Boudinot et al., 2008; Benichou et al., 2012).

ClonotypeR is the combination of support scripts to align and extract CDR3s, and a package for the R language for statistical computing, also available through Bioconductor.

## Approach

Detecting the V–J junction poses a challenge. The last nucleotides before the conserved cysteine are not enough to discriminate between every V segment, therefore it is necessary to sequence long reads. ClonotypeR was originally designed for the output of Roche 454 sequencers, where the error rate in the order of 4 % (Margulies et al., 2005), and most errors are insertions and deletions caused by homopolymers. As a consequence, the error rate will vary between V segments, which makes difficult to filter the alignments on quantitative quality scores. We therefore resorted to a two step strategy.

In the first step the V segments are detected by comparing the whole reads with a list of reference V segment sequences. ClonotypeR includes a list for mouse that was prepared using the NCBI Reference Sequence (Pruitt et al., 2007) entries NG_007044 (locus α and δ), NG_006980 (β) and NG_007033 (γ). The detection of V segments is done with the `bwasw` algorithm of BWA (Li and Durbin, 2010), that was designed for reads longer than 200 nt. This tool reports a mapping quality score (Li and Durbin, 2009) that represents the confidence that the alignment is unique and accurate. Since by definition the reads aligning to identical reference sequences have null mapping qualities, we removed redundant V segments from our reference list. BWA also reports also an alignment score that represents more directly the quantity of matches and mismatches. To avoid that a clonotype’s non-templated CDR3 would contribute to the alignment score, we trimmed the V reference segments after the conserved cysteins, which we identified by aligning all the V segments using SeaView (Gouy et al., 2010) in codon color mode. The final reference list combines segments from the α, β, γ and δ loci, therefore clonotypeR can detect accidental contaminations of one sample type with another during the preparation or the sequencing of the libraries, as well as rare Vα–Jδ rearrangements.

In the second step, the CDR3 boundaries are confirmed using the `vectorstrip` program from EMBOSS (Rice et al., 2000), requesting a perfect match with 20 nucleotides before the conserved cysteine. This independent detection of the V–CDR3 junction acts as a second filter for stringency and prevents from accidental frame shifts in case the BWA alignments did not reach the cysteine codon. The boundary of the J segment, downstream the FGxG motif, is then searched with the same strategy. The CDR3 is then translated using BioPerl (Stajich et al., 2002), and the data is saved in a tab-separated text file.

Using the Bioconductor package, the data table is loaded in R. Unproductive rearrangements that resulted in a frameshift or a stop codon are flagged on the fly. The data includes the mapping quality and alignment scores, so that their distribution can be studied in order to determine thresholds under which the low-quality clonotypes are discarded, if necessary. The analysis of the data (Figure 1) typically continues with differential representation analysis, sample clustering, etc., using standard R packages. ClonotypeR provides additional functions for operations that are not available in other packages, in particular for producing lists of clonotypes that are unique or common to groups of samples. Quantitative analysis can be done at multiple levels, for instance V or J segments, translated CDR3s, or combinations, in particular V, CDR3, J, which characterizes the clonotypes. The abundance of CDR3s can also be quantified using their nucleotide sequence.

**Figure 1.**
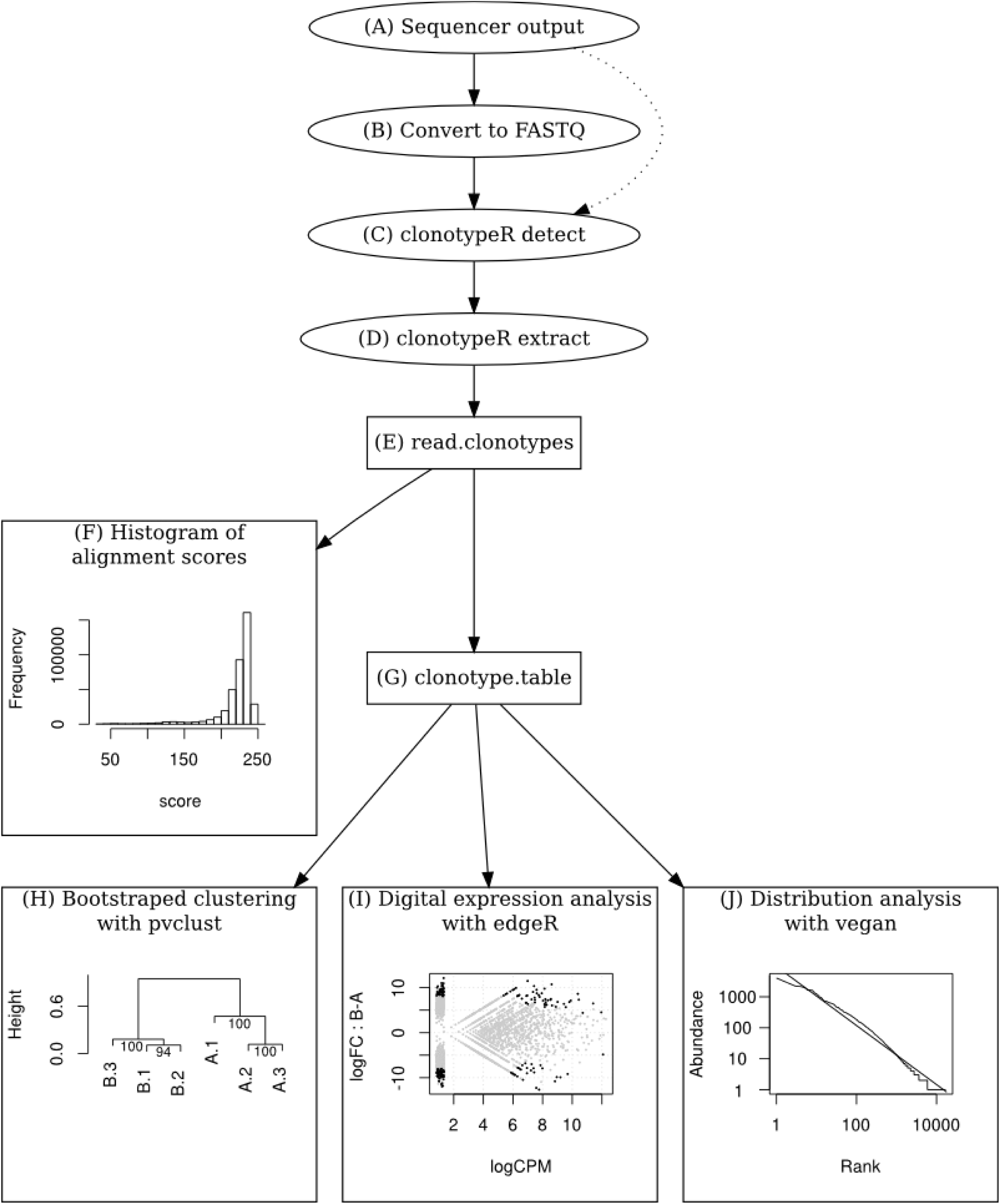
(A, B) Convert the sequences from SFF or AB1 format to FASTQ using `sff_extract` (http://bioinf.comav.upv.es/sff\_extract/download) and EMBOSS respectively. (C) Detect the V segments with the command `clonotypeR detect.` (D) Extract the clonotypes with the command `clonotypeR extract.` The output is a table indicating the V and J segments, alignment score and mapping quality, CDR3 sequence and quality. (E) In R, load the table using the `read_clonotypes` command. (F) Inspect the distribution of the alignment scores and the mapping qualities to identify thresholds below which outliers are discarded. The alignment scores vary with the length of the matching region between the reads and the reference segments, and therefore with the primer design for the cDNA amplification. (G) Produce a consolidated expression tables using the `clonotype_table` command. (H) Cluster the clonotype libraries, for instance with the pvclust R package. (I) Identify clonotypes enriched in groups of libraries by digital expression analysis, with packages such as edgeR. (J) Study the distribution of the CDR3 expression levels with the vegan package, to compare the repertoire structures and richness.

As a compact representation of nucleotide-characterized clonotypes, we implemented of the nomenclature of (Yassai et al., 2009), which assigns a unique synthetic identifier to each clonotype. However, tests on large scale data revealed that this nomenclature sometimes produces identifiers that are not unique to a clonotype. For example, aAn.1A14-1A43L9 represents a clonotype of 9 aminoacids, made by the junction of the TRAV 14-1 and TRAJ 43 segments, where the non-templated part is a GCT codon for alanin, inserted between a templated alanin from the V segment and a templated glutamin from the J segment. We show in Table 1 that this identifier corresponds to more than one clonotype. We propose to solve this problem by using a “long” variant of the identifier, where the full translated sequence is represented. Unfortunately, this long format would also have its own identifier collisions, as in the second example of Table 1: if the clonotype does not contain non-templated nucleotides, the long format would not indicate where the V–J junction is. We solve this problem by indicating the position of the junction by a number at the place where the identifier would contain codon IDs if there were non-templated nucleotides. To our knowledge, this “long” identifier format produces unique clonotypes, and is still reasonably short for usage as row identifier in R.

**Table 1.**
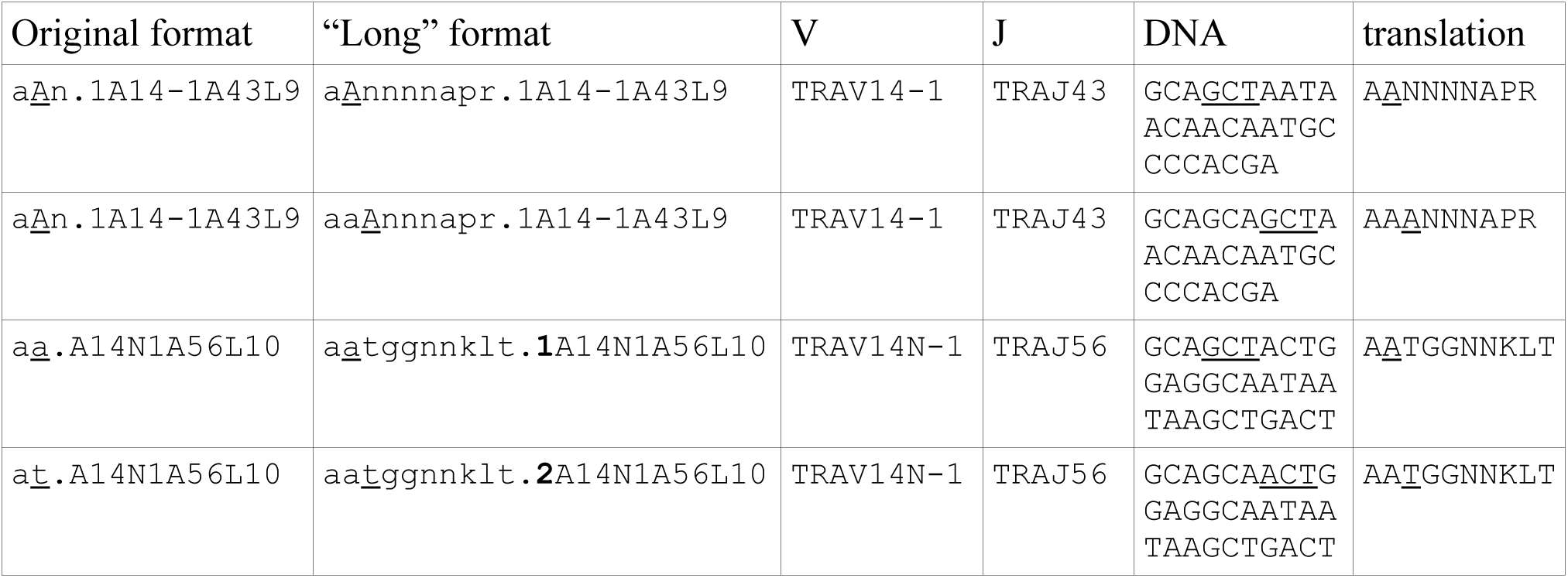
A “long” clonotype identifier based on the original format of Yassai et al., 2009. The first two rows show a clonotype that is be ambiguous in the original format but not in the long one. The second pair of rows shows two clonotypes that would be amiguous in the long format if the position of the V–J junction would not be indicated after the peptide sequence (bold). Critical differences between pairs of rows are underlined.

## Discussion

ClonotypeR differs from Decombinator (Thomas et al., 2013) by its design taking advantages of long reads. On-line tools like IMGT/HighV-QUEST (Alamyar et al., 2012) are limited in the amount of data they accept. Moreover, local processing gives full control of the software pipeline used, and therefore reproducibility of the results. Local installations in virtual machines are also becoming relevant for cloud computing.

ClonotypeR accepts user-provided reference alignments, for instance to analyze human data, take into account genetic polymorphisms, or study B cell receptors (BCRs). However, because of the somatic hypermutations, work on BCRs will need further adjustments.

To study colonotype abundance at the nucleotide level, we introduced a modification of the unique identifiers introduced by Yassai et al. (2009), in order to keep a strict one-to-one correspondence between clonotypes and their identifiers. This format is slightly longer, and might be shortened by removing the last part indicating the number of aminoacids, since in our modified version they are all represented. Alternatively, it may be interesting to replace this length information by a measurement of the edit distance between the clonotype and the juxtaposed V and J segments.

For our initial release, we focused on T cell receptors sequenced in Roche 454, Illumina, or capillary sequencers. However, ClonotypeR can handle the typical throughputs produced by HiSeq instruments. On a desktop computer running Debian Linux with a quad-core hyperthreaded CPU and 12 Gb of RAM, we could reprocess the 38 million CD8 ^+^ TRα (TRA) reads deposited as SRR407172 in NCBI GEO by (Genolet et al., 2012) in 10 hours. This deep data was obtained from naive mice and therefore broadly covers the 4,142 possible V–J segment combinations. Our results were in good match with the original analysis, with respectively 28 vs. 29 millions aligned reads, (17 vs. 15 millions unambiguous), and 3,653 vs. 3,901 different combinations of V and J segments.

ClonotypeR runs on Linux and Mac OS, as well as Windows (Bioconductor only). It fills the gap between the sequencer and the data analysis, without requiring online data submission, and facilitates the application of tools designed for digital gene analysis and diversity analysis to the field of immune repertoire studies.

## Acknowledgments

### Funding

We thank Jack Gorsky for useful comments on the Yassai nomenclature. This work was supported by research grants from MEXT and JSPS to SH (19059014 and 20689012), from MEXT to RIKEN CLST and RIKEN OSC, and from a RIKEN presidential grant. EMF received a JSPS long-term post-doctoral fellowship (P08627).

